# A single cell atlas of the developing *Drosophila* ovary identifies follicle stem cell progenitors

**DOI:** 10.1101/732479

**Authors:** Maija Slaidina, Torsten U. Banisch, Selena Gupta, Ruth Lehmann

**Affiliations:** HHMI, Skirball Institute of Biomolecular Medicine, Department of Cell Biology, NYU School of Medicine, NY, USA

## Abstract

Addressing the complexity of organogenesis at a system-wide level requires a complete understanding of adult cell types, their origin and precursor relationships. The *Drosophila* ovary has been a model to study how coordinated stem cell units, germline and somatic follicle stem cells, maintain and renew an organ. However, lack of cell-type specific tools have limited our ability to study the origin of individual cell types and stem cell units. Here, we use a single cell RNA sequencing approach to uncover all known cell types of the developing ovary, reveal transcriptional signatures, and identify cell type specific markers for lineage tracing. Our study identifies a novel cell type corresponding to the elusive follicle stem cell precursors and predicts sub-types of known cell types. Altogether, we reveal a previously unanticipated complexity of the developing ovary, and provide a comprehensive resource for the systematic analysis of ovary morphogenesis.

## Introduction

Organs are often maintained by tissue specific adult stem cells, which reside in specialized niches and contribute to tissue maintenance during the lifetime of the organism. These niche:stem cell compartments are established during development, and are tightly regulated during adulthood to ensure organ homeostasis in changing environmental conditions and during aging. Dissecting the origins and molecular mechanisms of adult stem cell specification and morphogenesis is challenging. In many systems, it is unclear whether adult stem cells are direct descendants of embryonic progenitors or whether they are specified later during development.

*Drosophila melanogaster* is a genetically tractable organism and their ovaries have served as a model for adult stem cell studies since decades. However, addressing cell type specific functions and how cells regulate each other to establish an adult organ has been hampered by lack of cell type specific tools and markers. Here, we report on a comprehensive single cell atlas of the developing *Drosophila* ovary and identify the progenitors of adult stem cell units. *Drosophila* ovaries house two adult stem cell units: germline stem cell (GSC) and follicle stem cell (FSC) (Dansereau and Lasko, 2008), thus providing an excellent model system to study adult stem cell development and regulation in a genetically tractable organism. The major ovary function, egg production, is achieved by coordinated proliferation and differentiation of GSCs and FSCs, which are both regulated by specialized somatic niche cells. The GSC daughter cells differentiate into eggs, while cells deriving from FSCs give rise to an essential follicle epithelium that covers and nurtures the egg and provides the developing oocyte with essential axial patterning information (Riechmann and Ephrussi, 2001). Numerous studies of GSCs have revealed key principles of niche:stem cell signaling, and delivered a wealth of knowledge of GSC development and establishment. On the other hand, the exact origin of FSCs remains elusive and their development has yet to be studied. In addition to GSCs and FSCs, ovaries contain a number of other somatic cell types that support the development and adult functions of the ovary. During development, their proliferation, movement and differentiation needs to be coordinated to establish a functional adult organ. How this is orchestrated and the exact function of individual cell types remains to be elucidated. This knowledge gap is partly caused by a lack of cell type specific markers and tools.

Single cell RNA sequencing (scRNA-seq) technology allows to capture transcriptomes of individual cells of an entire organ (Stuart and Satija, 2019). We applied this technology to developing fly ovaries to gain a systems view of the complete repertoire of ovarian cell types and their functions during development. For our studies, we chose the late third larval instar (LL3) stage for two reasons. First, specific progenitor populations for the majority of cell types are thought to be established by this stage and, second, germ cells transit from primordial germ cells, which remain undifferentiated and proliferate, to self-renewing stem cells that reside adjacent to their somatic niches and produce more proximally located differentiating cysts, which will give rise to the eggs (Gilboa, 2015).

Utilizing scRNA-seq, we identified all known ovarian cell types, additional sub-types of these cell types and a novel cell type. By lineage tracing and genetic cell ablation experiments, we demonstrated that this novel cell type corresponds to the long sought after FSC progenitors. Furthermore, we computed transcriptional signatures for all cell types in the developing ovary, started predicting their function using Gene List Annotation tools, and selected cell type-specific markers that can be used for further interrogation of cell type function and lineage tracing. Our work provides a resource for future morphogenesis studies of niche:stem cell unit establishment and gonadal support cell function.

## Results

### Single cell RNA sequencing of developing *Drosophila* ovaries

For single cell RNA sequencing (scRNA-seq) analysis, we dissected ovaries from developing larvae at LL3 stage that expressed a His2AV::GFP transgene. In these animals, all cell nuclei were labeled with GFP (Figure S1A), allowing cell purification from debris by fluorescence-activated cell sorting (FACS) (Figure 1B). scRNA-seq was performed on two independently collected samples using the 10x Genomics Chromium system for complementary DNA (cDNA) conversion and amplification, library preparation, and sequencing. We obtained 753 and 1,178 single cell transcriptomes from approximately 15 and 45 larval ovaries, respectively, and used Seurat v2 (Butler et al., 2018; Satija et al., 2015) pipeline to perform established quality control (QC) steps. By plotting the number of genes detected per cell transcriptome, we uncovered two distinct cell populations segregating by the number of genes detected (Figure S1B). Subsequent analyses using known germ cell marker genes (including, *vas*, *AGO3* and others) determined that the population with higher number of genes detected are germ cells (4930±36 in germ cells vs. 2931±17 in somatic cells (mean±SEM)) (see Supplemental Note and Methods) (Figure 1C, S1C). Moreover, we detected also a higher number of unique molecular identifiers (UMIs) in germ cells than in somatic cells (53,531±1001 vs. 21,097±27) (Figure 1C, S1D), suggesting that germ cell contain higher RNA levels than somatic cells. Therefore, we manually separated germ cell transcripts from somatic cell transcripts for initial QC steps (Supplemental Text and Methods). Subsequently, we retained 699 and 1,048 high quality cell transcriptomes from the two samples, respectively. Gene expression levels highly correlated between both replicates (Spearman = 0.97, Figure 1D), and between our scRNA-seq dataset and bulk RNA-seq generated from dissected LL3 ovaries (Spearman = 0.87, Figure 1E) despite different library preparation methods (see Methods). Thus, our sample preparation methods are robust and did not significantly alter ovarian transcription profiles. Together, scRNA-seq of dissected developing ovaries yielded a high-quality dataset containing 1,747 ovarian cell transcriptomes.

**Figure 1.**
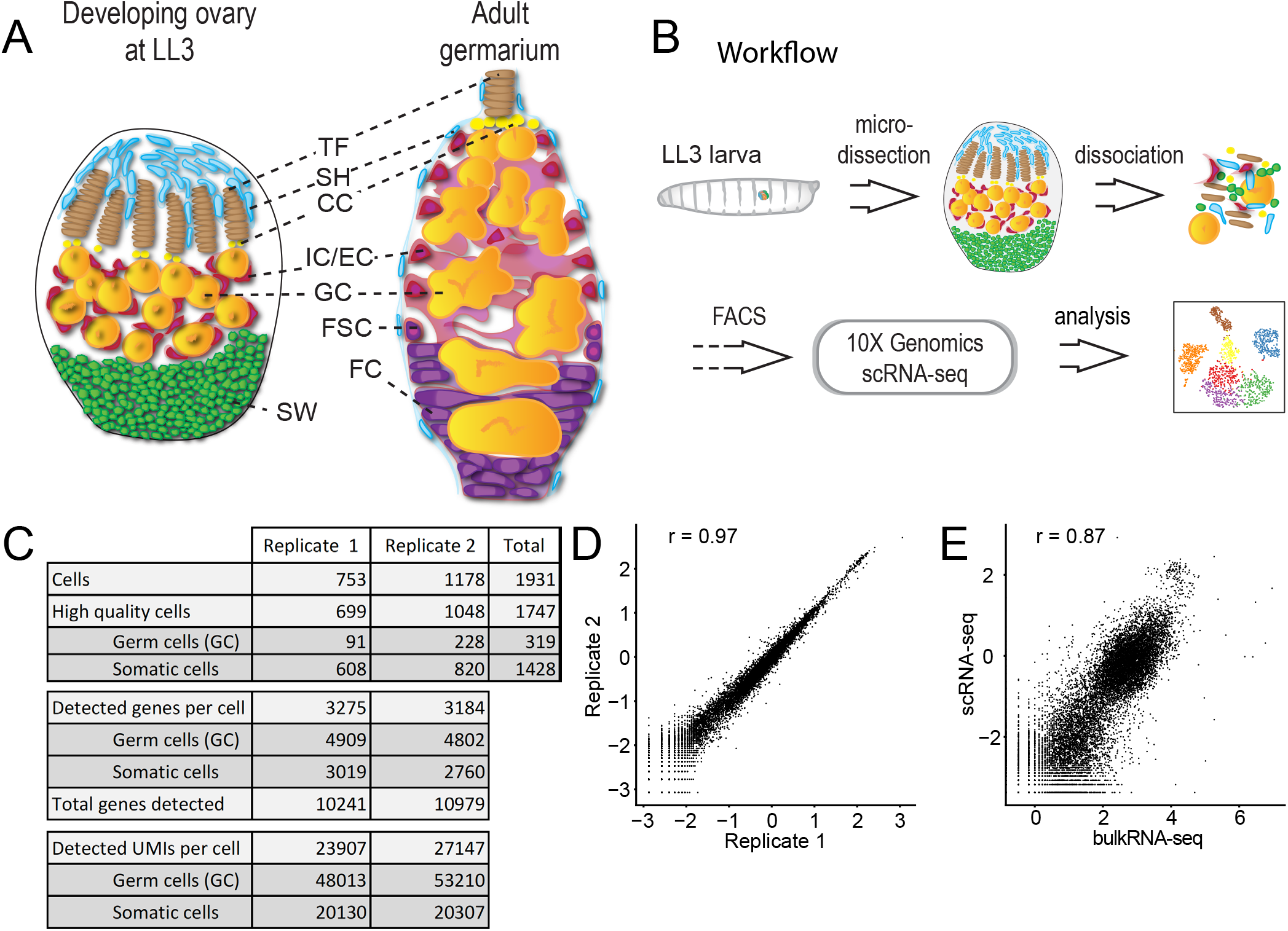
scRNA-seq experiment design and statistics. A – schematic drawing of a developing ovary, and adult germarium. B – scRNA-seq experiment workflow. C – scRNA-seq experiment statistics. D – gene expression averaged among individual cells in each replicate and compared to each other. E – gene expression in replicate 1 averaged among individual cells and compared to bulk RNA-seq.

### The cell types of the developing ovary

Next, we determined the cell type identity for each high-quality cell transcriptome. We aligned the datasets from the two independent experiments, reduced dimensionality and binned cells into clusters using unsupervised hierarchical clustering (Butler et al., 2018; Satija et al., 2015) (Figure 2A). With multiple clustering parameters, we robustly identified seven clusters (Figure 2A, S2AB). Previous studies had identified six cell types in the larval ovary based on morphology, position and select gene expression (Figure 1A): germ cells (GCs) located in the middle of the ovary and five somatic cell populations surrounding the germ cells. Sheath cells (SH) are located at the anterior tip of the LL3 ovary (King et al., 1968). During metamorphosis they will subdivide the ovary into 16-20 units, called, ovarioles (Irizarry and Stathopoulos, 2015; King et al., 1968). Terminal filaments (TFs) and cap cells (CCs) together form the niche for GSCs (Gilboa, 2015; Sahut-Barnola et al., 1996; Song et al., 2007) and are positioned between the SH and GCs. Intermingled cells (ICs) have acquired their name because they intermingle with germ cells and regulate their proliferation(Gilboa and Lehmann, 2006; Li et al., 2003). Finally, swarm cells (SW) (also called basal cells) are located at the posterior tip of the LL3 ovary and their function is not known (Couderc et al., 2002; Gilboa, 2015).

**Figure 2.**
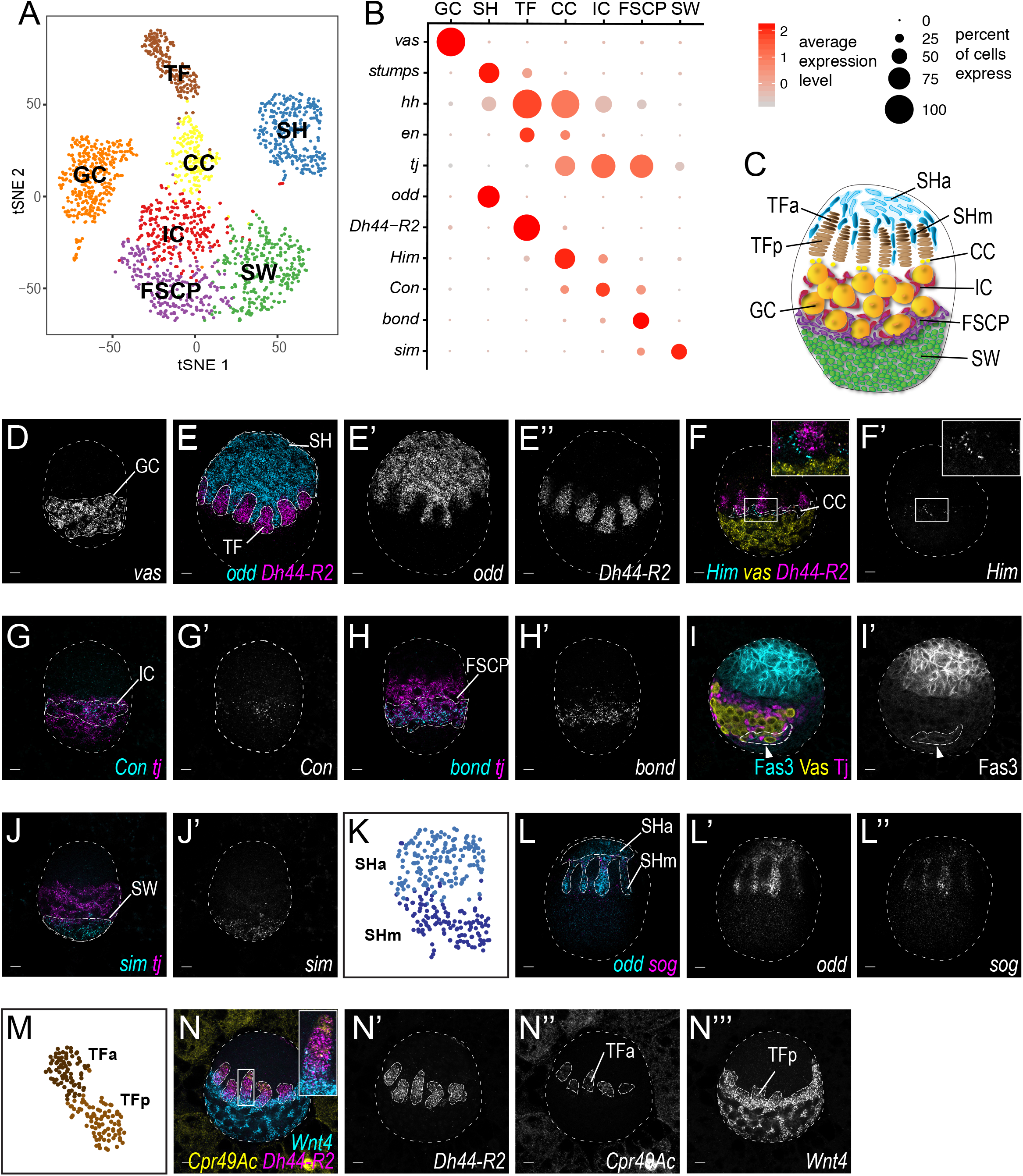

To correlate the clusters obtained through scRNA-seq with previously described cell types, we identified markers for each cluster based on a) enrichment for the cluster relative to other clusters, b) robust level of expression, and c) expression in a large fraction of cells within a given cluster (Supplemental Table 1). We then compared these markers to previously described cell type marker genes (Figure 2B, S2C). A germ cell cluster was easily identified by known germ cell specific genes, such as *vas* (Figure 2BD, S2G). In contrast, assignment of somatic cell clusters to specific cell types was more difficult as many genes are expressed by several cell clusters. Nonetheless, we were able to assign SH fate based on *stumps* and *ths* expression (Irizarry and Stathopoulos, 2015), TF and CC fates on the basis of high *hh* expression in both (Lai et al., 2017) and alternate enrichment for *en* in TFs (Forbes et al., 1996) and *tj* in CC (Li et al., 2003) (Figure 2B, S2CI-M). We confirmed these cell type assignments by assessing the expression patterns of newly identified cluster specific markers using a highly sensitive method for *in situ* visualization of RNA, hybridization chain reaction (HCR) (Choi et al., 2018). These analyses revealed that *odd* labels SH (Figure 2E, S2J), *Dh44-R2* labels TFs (Figure 2EF, S2N), and *Him* labels a narrow band of CCs flanked by TF and IC cells, respectively (Figure 2F, S2O), thus confirming our initial cell type assignments.

To assign identities to additional clusters, we identified novel cluster-specific markers (Figure 2B, Supplemental Table 1). Two *tj*-expressing clusters remained to be assigned. Since ICs and their presumed adult descendants escort cells (ECs) and adult FSCs/FC express *tj* (Gilboa and Lehmann, 2006; Li et al., 2003), we speculated that these clusters could correspond to ICs and the elusive FSC progenitors (FSCPs). Two markers, *Con* and *bond* (Figure 2BGH, S2PQ) were differentially expressed between the two clusters, and by anatomical position, we determined that *Con*-expressing cells are intermingled cells (IC) (Figure 2GH). We hypothesized that *bond*-expressing cells, which reside posterior to ICs and are in direct contact with germ cells, are putative FSC progenitors (FSCP). In support, these putative FSCPs also expressed *Fas3* (Figure 2I, S2CR), which labels follicle cells and FSCs in adults (Nystul and Spradling, 2007). Lastly, *sim* was specifically expressed at the posterior tip of the ovary identifying SW (Figure 2BJ, S2S). Thus, by correlating known and newly identified markers with expression patterns in the developing ovary, we were able to assign seven clusters to distinct ovarian cells types and identified a putative progenitor population for the adult FSCs. Moreover, we have obtained gene expression profiles of all the cells types in developing ovaries (Supplemental Table 2).

### Transcriptionally distinct sub-types divide TF and SH cells

After assigning each cluster with a cell type identity, we searched for systematic transcriptome variability within clusters. For this, we raised the resolution parameters for *in silico* cell clustering, and the GC, TF and SH clusters split into sub-clusters (Butler et al., 2018) (Figure S2B, arrowheads, 2KM). Further investigation suggested that the GC cluster split is unlikely to have biological significance as it was not observed when the cluster was analyzed separate from the somatic cell types (Figure S2D, and Supplemental Text).

To test the robustness of TF and SH sub-clusters, we re-clustered each cell type independently of other cell types (Figure S2EF). The gene expression patterns of the independently re-clustered SH and TF sub-clusters clearly corresponded to the initially identified clusters (Spearman = 0.99). Thus, SH and TF sub-clusters may represent specific sub-populations among SH and TF cells. To determine whether these sub-populations reflect a developmental or morphological distinction within the known cell type, we identified markers that distinguished the sub-clusters (Supplemental Table 1) and assessed their expression patterns *in vivo*. For the SH sub-clusters, we found that SH cells expressing both *sog* and *odd* are migrating between the TF stacks, hereafter referred to as SHm (migrating) (Figure 2L, S2TU), and that SH cells, which only express *odd* correspond to the SH cell located at the anterior tip of the ovary, which we now call SHa (anterior). For TF subtypes, *Dh44-R2* labeled both TF sub-clusters, while *Cpr4Ac* was expressed only in the anterior half and *Wnt4* labeled the posterior half of the TFs as well as other cell types (CC, IC, FSCPs and SW) (Figure 2N, S2V-X). We refer to these subtypes as TFa and TFp for anterior and posterior, respectively.

### Cell type specific transcriptional signatures reveal functional connections between cell types

Cell states and functions should be reflected by the gene repertoire they express. Thus, the transcriptomes for each cell type in developing ovaries should allow us to explore their respective functions. To enrich for transcriptional signatures that are cell type specific, we first excluded those genes that are uniformly expressed between cell types and mostly encode proteins associated with general cellular processes (Supplemental Table 1). Next, we excluded marker genes that were assigned to more than three cell types. We visualized the gene expression levels of these transcriptional signature in each cell type by heatmap (Figure 3A and 3B, Supplemental Table 3). Finally, we exploited the Gene List Annotations for *Drosophila* (GLAD) online resource (Hu et al., 2015), hypergeometric tests and manual curation to correlate transcriptional signatures with potential functional specializations for each cell type (Figure S3A, Supplemental Table 3).

**Figure 3.**
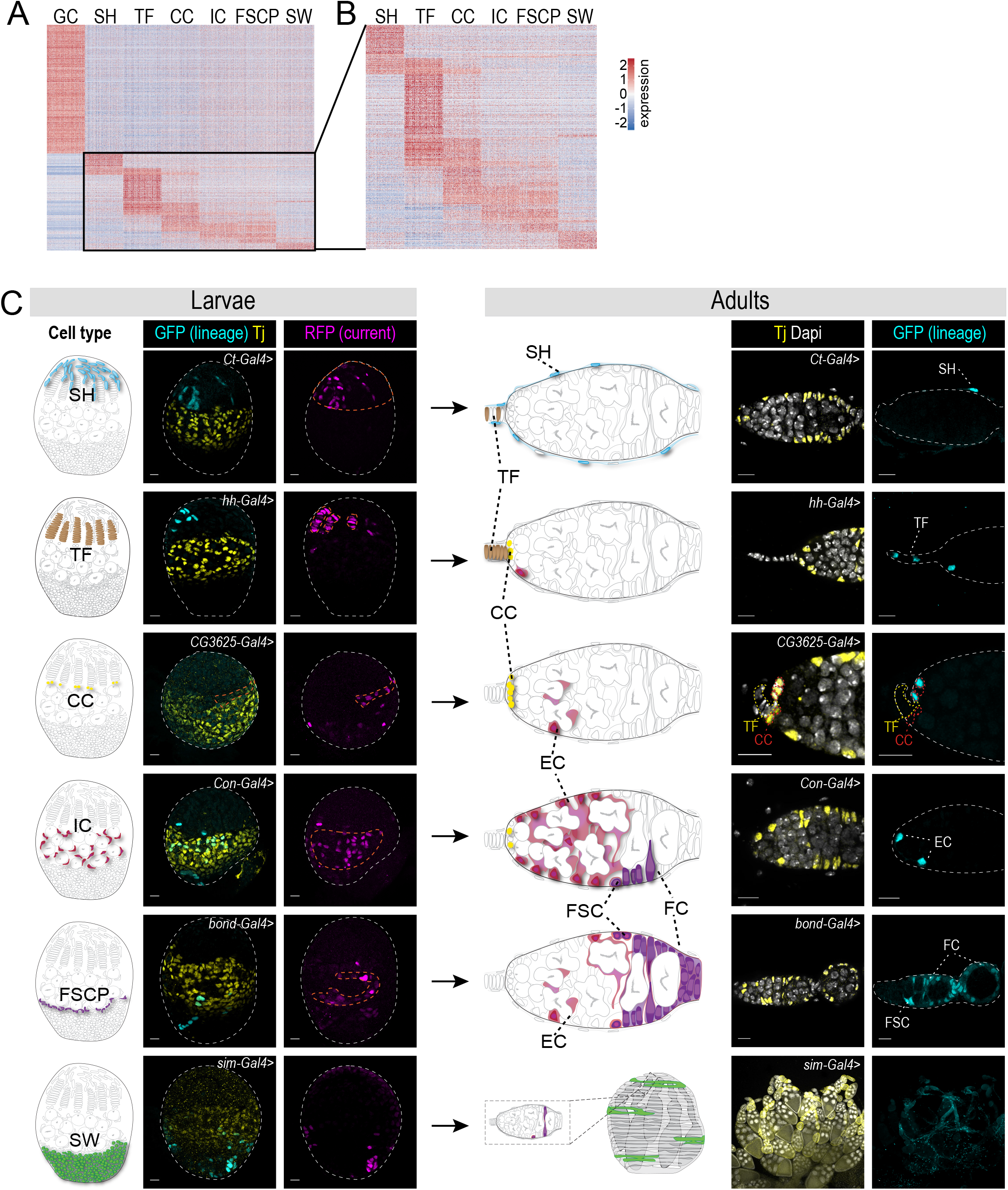

Germ cells are specified earlier and independently of the somatic cells of the gonad. Consistently, germ cells have the highest number of signature genes (1073 genes) (Figure 3A). The germ cell transcriptional signature was enriched for multiple GLAD categories that fit well with known and prominent features of the germline, including posttranscriptional regulation (splicing, RNA regulation, translation, protein degradation) and mitochondrial maturation and selection (oxidative phosphorylation, autophagy) (Cox and Spradling, 2003; Kai et al., 2005; Lieber et al., 2019; Slaidina and Lehmann, 2014; Teixeira et al., 2015).

All somatic cells of the gonad arise from a somatic gonadal precursor population that is specified during embryogenesis (Boyle and DiNardo, 1995; Moore et al., 1998; Riechmann et al., 1998). The signature gene heatmap reflects this shared origin as most cell types are identified by only a small number of genes. We arranged the somatic gonadal cell types along the ovarian anterior-to-posterior axis in a pattern that may reveal developmental relationships between cell types and distinct functional specialization (Figure 3B). For example, the two most anterior cell types, SH and TFs have clear transcriptional signatures that set them apart from each other and all other somatic gonadal cell types (SH – 179 and TF – 404 genes), suggesting that these cell types may have diverged from a common precursor pool early in development (Godt and Laski, 1995). Consistent with their common function in forming the germline stem cell niche, CCs and TFs share a fraction of their transcriptional signature (CC – 262, common – 122 genes). Finally, the strongly correlated transcriptional signature of ICs and FSCPs (IC – 155, FSCP – 161, common – 101 genes) suggests that they originated from a common progenitor subpopulation late in development or that they fulfill related functions.

Consistent with their role in germ cell support and gonad morphogenesis, all somatic cell types are enriched for gene ontology terms associated to **cell signaling**. For example, Notch ligands are enriched in SH and TFs, and all other cell types highly express Notch receptors and/or their downstream pathway components. Multiple somatic cell types are significantly enriched for genes expressing proteins with roles in mediating **cell-cell communication**. These protein classes will expand previous observations which showed that, in addition to conventional signaling pathways, ovarian cells coordinate their behaviors through alternative modes of signaling (Banisch et al., 2017; Gilboa et al., 2003). For example, a gap junction protein *zpg* is highly enriched in germ cells, while other gap junction proteins, *inx2*, *inx3*, and *ogre,* are expressed in somatic cell types surrounding the germ cells. Cell-type specific enrichment for **transcription factors and DNA binding proteins** can provide a useful tool to study gene regulatory networks for gonad development at a cell type resolution. For example, the *bab1* and *bab2* transcription factors are required for the development of TFs and CCs, the GSC niche (Godt and Laski, 1995). Adult FSCs are regulated by the Hedgehog signaling pathway (Zhang and Kalderon, 2001), and its downstream transcription factor *ci* is specifically enriched in FSCPs suggesting that similar regulation might take place during development as well. For a more detailed analysis and summary of gene classes enriched in specific cell types, refer to Figure S3A and Supplemental Table 3.

### Connecting precursors to adult cell types by lineage tracing

During metamorphosis, developing ovaries turn into the adult structure. Yet, for only a fraction of adult descendants are the progenitor cell types known. Therefore, we took advantage of our newly identified cell type markers and designed lineage tracing experiments that determined the lineage relationships between defined cell types in developing ovaries and adult descendants. For lineage-analysis, we used G-TRACE, a method that combines a Gal4 driver with the UAS FLP recombinase-FRT systems to generate clones for lineage analysis and a UAS- fluorescent reporter to observe real-time expression patterns in combination with cell type-specific Gal4 driver (Evans et al., 2009). To identify appropriate driver lines for each cell type, we tested 79 publicly available lines with *Gal4* integrated near the regulatory sequences of individual somatic cell type marker genes. We first tested the expression pattern of each line at LL3 by co-staining with anti-Tj antibodies and Dapi. We identified at least one Gal4 driver line for each somatic cell type (Figure 3C, S3BC). While these drivers were predominantly expressed in the desired cell types, they were also labeling a few cells of other cell types, suggesting expression in a common progenitors (Figure 3C, S3C). Real-time labeling with the G-TRACE cassette showed relatively sparse expression, but allowed us to efficiently mark cell lineages and determine the cell type lineage relationships between larval and adult ovaries (Figure 3C, S3BC). For example, *cut*-Gal4 labeled SH cells at LL3 gave rise to the epithelial sheath that surrounds each ovariole in the adult (Figure 3C, S3C) (Irizarry and Stathopoulos, 2015). As described previously, *hh-Gal4* marked the TF continuously from the larva to the adult (Figure 3C, S3C) (Lai et al., 2017). A *CG3625-Gal4* line predominantly labeled CCs at LL3 (Figure 3C, S3C, S4B) and their progeny gave rise to adult CCs and rarely ECs, which are thought to be IC descendants, suggesting common ancestry of both cell types (Song et al., 2007). Consistently, ICs labeled by *Con*-Gal4 gave rise to ECs and less frequently to CCs in adults (Figure 3C, S3C). The adult descendants of SW cells had not been determined. A *sim*-Gal4 driver labeled SW at LL3 (Figure 3C), however we did not detect any robust lineage expression in the adult germarium (besides rare labeling of single SH, EC and FCs, which might be due to a broader expression of *sim* at LL3). Instead, we identified lineage-labeled cells in the outer ovarian sheath (also called peritoneal sheath, (Spradling, 1993)) suggesting that it originates at least in part from the larval SW population (Figure 3C, S3C).

Finally, we chose *bond-Gal4* to analyze the putative FSCP population. We found that cells expressing this marker in the larva gave rise predominantly to follicle cells and FSCs in the adult (Figure 3C, S4C). We noted that *bond-Gal4* drives expression in a slightly broader pattern than what was observed with an RNA probe for the *bond* gene in the LL3 ovary (Figure 2H, 4C), this may explain sparse lineage expression in ECs and some other cell types in the adult (Figure 3C). Together, these results suggest that follicle cells are derived from a larval precursor population nested between the ICs, the precursors of the adult ECs, and the SWs.

**Figure 4.**
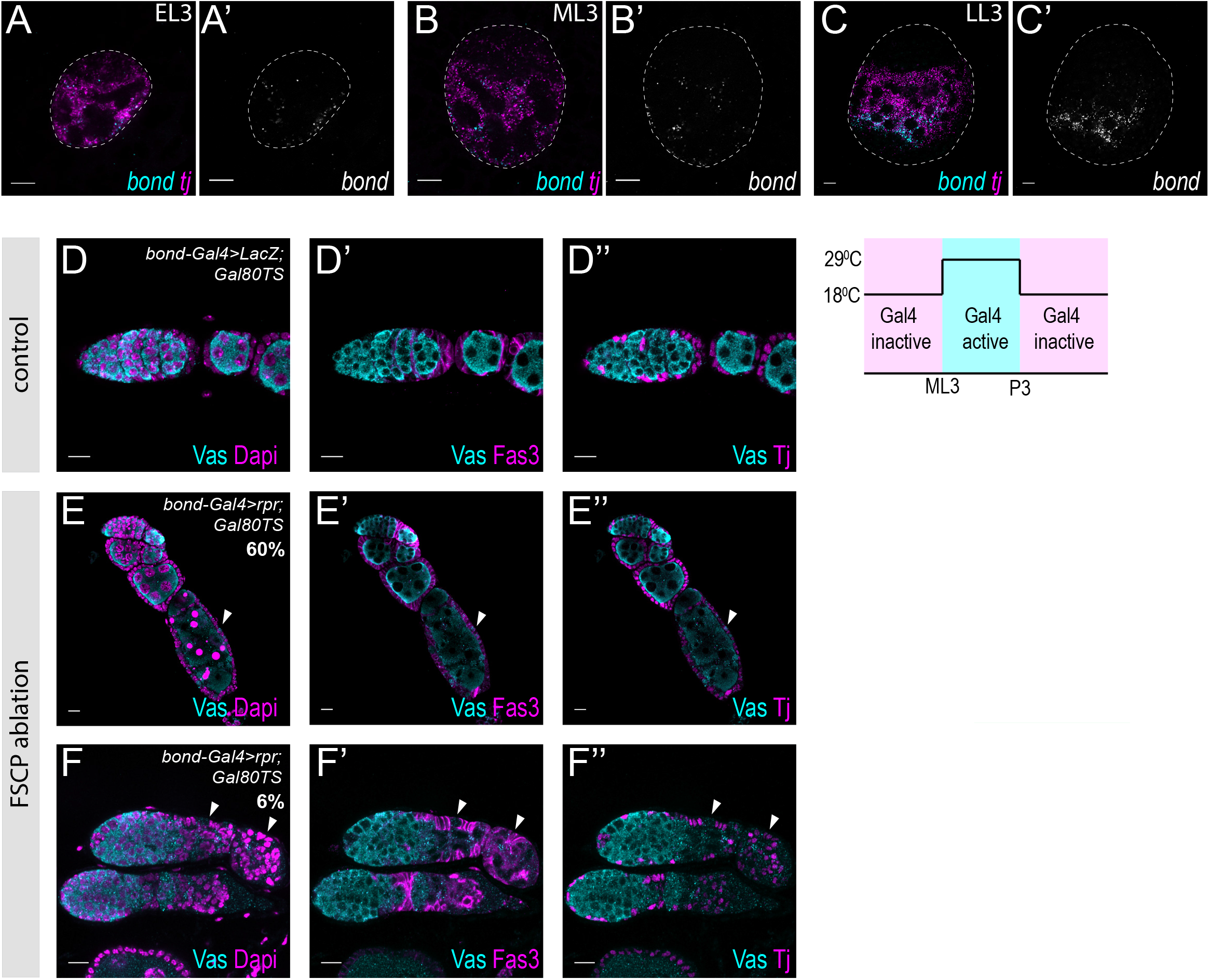
FSCPs give rise to FSCs and FCs in adult ovaries. A-C - mRNA in situ hybridization using HCR at EL3 (A), ML3 (B), and LL3 (C). tj (magenta) and bond (cyan). Scale bars - 10 m. D-F - Ablation of FSCPs by bond-Gal4 driven expression of the pro-apoptotic factor rpr restricted to late larval and early pupal stages causes FC defects in adults. D – control; E, F- predominant phenotypes observed upon rpr expression using bond-Gal4. E - follicular epithelium is distorted and germ cells are dying (arrowhead). F - follicle cells are distorted and no egg chambers are formed (arrowhead). Vas (cyan) labels GCs, Dapi (magenta in D-F) labels DNA, Fas3 (magenta in D’-F’) labels FSCs and FCs, Tj (magenta in D”-F”) labels most somatic cells of the ovary, including escort cells and follicle cells. Gal4 activation (290C) and inactivation (180C) by Gal80TS is indicated in a schematic drawing on the right. Note - in 4F, germ cell number in germarium is slightly increased, which might arise due to distorted EC functions and subsequent block in GC differentiation. Scale bars - 20 um.

### FSCP ablation during development causes follicle cell defects in adults

After identifying a FSCP population that gives rise to adult FSCs and follicle cells, we asked when these cells are first specified by observing the expression of *bond* RNA during earlier stages of development. We detected *bond* expression first at the early-third larval instar stage (EL3, 72h AEL (after egg laying)) (Figure 4A, S4D), and more robustly at mid-third larval instar (ML3, 96h AEL) (Figure 4B, S4E). At EL3, sparse and weak *bond* expression covers the entire *tj* expression domain, suggesting that the FSCPs share common ancestry with ICs, which also express *tj*. At ML3, *bond* starts to enrich more posteriorly, and a strong band of *bond* expression is present at the posterior part of the *tj* expression domain at LL3, now likely restricted to the FSCP lineage (Figure 4C, S4F).

To test whether FSCs and follicle cells arise from this larval cell population, we ablated the *bond-Gal4* expressing cells specifically from ML3 to early-pupal stages (144h) using Gal4/Gal80TS mediated temporal expression of *reaper* (*rpr)*, an apoptosis inducing gene (White et al., 1996). Control adult females experiencing equivalent temperature shifts, developed normal ovaries (Figure 4D). In *rpr*-expressing females, ∼60% of ovarioles showed defects in the follicular epithelium and dying germ cells, while other germarial structures – TFs, CC, GSCs, and ECs were unaffected, demonstrating the specificity of this defect to follicle cells (Figure 4E). In ∼6% of ovarioles, follicle cells were severely distorted and egg development was highly abnormal (Figure 4F). Altogether, these results indicate, that the *bond*-expressing precursor population marks FSCPs which predominantly give rise to the FSCs of the adult.

## Discussion

The development of *Drosophila* ovaries has been studied for decades. Nevertheless, functional studies of most ovarian cell types have been hindered by a lack of cell type specific markers and driver lines. Our study has identified cell type specific marker genes, and for over a hundred of these genes publicly available GFP fusion constructs exist (Sarov et al., 2016) that could be used for cell labeling and/or live imaging (Figure S4GH). We used a number of publicly available drivers for lineage analyses. While some of these were expressed in a broader pattern than expected from mRNA expression patterns, we were able to identify GAL4 driver lines for lineage tracing of each larval ovary cell type. In particular, these lines helped us to determine the adult descendants of the swarm cells and identified the long sought-after follicle stem cell progenitors. Going forward, the cell-type specific markers identified in this study can be used for further tool building to more specifically and completely target individual cell types (Figure S4I). Furthermore, strategies involving Gal80TS and split-Gal4 systems can be used to improve driver specificity, and avoid expression in other tissues (Pfeiffer et al., 2010). Our GLAD analysis grouped cell type signature genes according to their molecular and cellular functions (Hu et al., 2015). Predicted cellular functions and protein classes enriched in each cell type will provide new insight into how cells in the developing ovary interact, how stem cell units are established and how these precursor cell interactions support the morphogenesis of the adult ovary.

A major finding of our study is the identification of a follicle stem cell progenitor population. Our results show that the transcriptional signatures of FSCPs and ICs are similar. This could indicate that these two cell types are specified from a common progenitor. In support, the FSCP marker gene *bond* is detected as early as EL3 in a broad expression domain spanning both the FSCP and IC progenitors. *bond* may be expressed in the common progenitor pool and later become restricted to the FSCPs, or the *bond*-expressing FSCPs may be initially dispersed and later migrate posteriorly. In addition to common developmental origins, an overlap in transcriptional signatures may also reflect shared functions. Consistently, ICs and FCs both intimately interact with germ cells and guide their differentiation (Banisch et al., 2017; Wu et al., 2008; Xie, 2013); thus, analyzing the overlap between the IC and FSCP transcriptional signatures might reveal the nature of IC/FSCP to GC signaling, and shed light on stem cell to support cell communication in general.

Altogether, our study provides a systems-wide overview of cell types, and their transcriptional profiles and signatures in the developing *Drosophila* ovary. This resource will facilitate future studies leading to a better understanding of how stem cell populations are specified, regulated and maintained in the context of a growing organ, and, more general, how a complex interplay of several cell types achieves to build an organ. Future scRNA-seq experiments using additional stages of development (earlier larval, pupal, adult) or using scRNA-seq methods that allow simultaneous lineage tracing, like scGESTALT (Raj et al., 2018) will allow to identify the complete lineage relationships between the ovarian cell types. Moreover, perturbing functions of individual cell types, will provide information about cellular processes that are coordinated between the cells and how this coordination is achieved. Together our work should provide an invaluable resource for the stem cell and developmental biology research communities.

## Acknowledgements

We thank Drs. Michael Buszczak, Dorothea Godt, Erika Bach for sharing reagents. We are grateful to Drs. Brian Oliver, NIH and Mark Van Doren, JHU for communication prior to publication. Transgenic fly stocks were obtained from the **Vienna Drosophila Resource Center (VDRC**, www.vdrc.at), the Bloomington Drosophila Stock Center (NIH P40OD018537) and KYOTO Stock Center (DGRC) at Kyoto Institute of Technology. The Fas3 antibody developed by Goodman, C. was obtained from the Developmental Studies Hybridoma Bank, created by the NICHD of the NIH and maintained at The University of Iowa, Department of Biology, Iowa City, IA 52242. We would like to thank the Genome Technology Center (GTC) for expert library preparation and sequencing, and the Applied Bioinformatics Laboratories (ABL) for providing bioinformatics support at the initial steps of the project. GTC and ABL are shared resources partially supported by the Cancer Center Support Grant P30CA016087 at the Laura and Isaac Perlmutter Cancer Center. Cell sorting technologies were provided by NYU Langone’s Cytometry and Cell Sorting Laboratory, which is supported in part by grant P30CA016087 from the National Institutes of Health/National Cancer Institute. We are grateful to Claudia Skok Gibbs for assistance with statistical and computational analyses. M.S. was a Howard Hughes Medical Institute Fellow of the Life Sciences Research Foundation. S.G. is supported by Dean’s Undergraduate Research Fund Grant, R.L. is supported by the Simons Foundation, NIH R37HD41900 and is HHMI investigator.

## Materials and Methods

### CONTACT FOR REAGENT AND RESOURCE SHARING

Further information and requests for resources and reagents should be directed to and will be fulfilled by the Lead Contact, Ruth Lehmann.

### EXPERIMENTAL MODEL AND SUBJECT DETAILS

#### Fly husbandry

Flies were raised on medium containing yeast, molasses and cornmeal, and kept at 25 °C. The lineage tracing and ablation experiments were performed at 18 °C and 29 °C as indicated in text.

### METHOD DETAILS

#### Dissections

For EL3, ML3 and LL3, properly staged larvae were rinsed in PBS (for immunofluorescence) or DPBS (for RNA *in situ* hybridization) and sexed (if possible). Posterior part of the cuticle was removed using forceps, and specimens were inverted. Intestines were gently removed, leaving the fat body and other organs intact and attached to the cuticle.

For L2, properly staged larvae were rinsed and their anterior was removed, leaving most organs partly extruding from the cuticle.

Female adults were fattened on yeast for 2-3 days. Abdomens were removed using forceps and parts of intestine were removed, leaving ovaries partly covered by abdominal cuticle.

#### Immunofluorescence

All steps were done with gentle rotation. Specimens were fixed in PBS, Triton-X (Tx) 0.3%, Paraformaldehyde 4% for 20 minutes at room temperature (RT) with gentle rotation, washed twice with PBS, Tx 1%, blocked/permeated for 2 hours in PBS Tx 1%, normal goat serum (NGS) 5% at RT for 2 hours. Primary antibody was diluted in PBS Tx 0.3%, NGS 5% and incubated for 2h at RT or 4 °C overnight. Subsequently, specimens were washed in PBS, Tx 0.3% three times for 20 minutes in RT and in PBS, Tx 0.3%, NGS 5% twice for 30 minutes. Secondary antibodies and Dapi were diluted in PBS Tx 0.3% NGS 5% and incubated for 2h at RT or 4 °C overnight. Subsequently, specimens were washed in PBS, Tx 0.3% four times for 20 minutes in RT. Finally, specimens were equilibrated in Vectashield mounting medium at 4 °C overnight and pieces of larval fat body containing ovaries/adult ovarioles were mounted in Vectashield.

#### RNA in situ hybridization

All steps are done in using RNAse free reagents and supplies with gentle rotation, except for steps at 37°C. The protocol was adapted from Choi et al. (Choi et al., 2018). Specimens were fixed in PBS, Tween (Tw) 0.1%, Paraformaldehyde 4% for 20 minutes at RT, washed twice with PBS, Tw 0.1% at RT, dehydrated with sequential washes with 25%, 50%, 75% and 100% methanol in PBS on ice 5 minutes each. Samples were stored at −20°C at least overnight (up to one week). Samples were rehydrated with sequential washes with 100%, 75%, 50% and 25% methanol in PBS on ice, permeated for 2 hours in PBS Tx 1% at RT, post-fixed in PBS, Tw 0.1%, Paraformaldehyde 4% for 20 minutes at RT, washed twice with PBS, Tw 0.1% for 5 minutes on ice, washed with 50% PBS, Tw 0.1%/ 50% 5xSSCT (5xSSC, Tween 0.1%) for 5 minutes on ice, washed twice with 5xSSCT for 5 minutes on ice, incubated in probe hybridization buffer for 5 minutes on ice, pre-hybridized in probe hybridization buffer for 30 minutes at 37 °C, and hybridized overnight (16 – 24 hours) at 37°C. Probe concentrations were determined empirically, and ranged 4 −8 pmol of each probe in 1 ml, probe solution was prepared by adding probes to pre-warmed probe hybridization solution. After hybridization, specimens were washed 4 times with probe wash buffer for 15 minutes each at 37°C, and twice with 5xSSCT for 5 minutes each at RT. Specimens were equilibrated in amplification buffer for 5 minutes at RT. Hairpin solutions were prepared by heating 30 pmol of each hairpin for 90 seconds at 95°C and cooling at RT in a the dark for 30 minutes, and subsequently adding the snap-cooled hairpins to 500 μl of amplification buffer at RT. Specimens were incubated in hairpin solution overnight (∼16 hours) at RT, and washed multiple times with 5xSSCT – twice for 5 minutes, twice for 30 minutes and once for 5 minutes. Dapi was added in the first 30-minute wash. Specimens where equilibrated in Vectashield overnight at 4oC and mounted in Vectashield, or further stained using the Immunofluorescence protocol (see above).

#### Imaging

Imaging was performed using Zeiss LSM 800 and Zeiss LSM 780 confocal microscopes using 40x oil NA 1.3 objectives.

#### Ovary dissociation

15 – 50 LL3 ovaries were dissected per sample in ice-cold DPBS; majority of associated fat body was removed with forceps and dissection needles. For dissociation, ovaries were transferred to 9-well glass plates and incubated in dissociation solution (0.5% Type I Collagenase, 1% Trypsin 1:250 in DPBS) for 15 minutes with gentle rotation. The suspension was vigorously pipetted multiple times during the dissociation to enhance the dissociation efficiency. Enzymatic dissociation was stopped by adding Schneider cell culture media with fetal bovine serum (S-FBS). Starting from this step, all plastic materials – pipet tips, tubes, filters – were coated with S-FBS. Cell suspension was filtered through a custom-made 40micron cell strainer. The strainer was built by securing nylon mesh in a cap of a 0.2 ml PCR tube and cutting the bottom of the tube and the cap. Upon filtering, dissociated cells were purified by fluorescence activated cell sorting (FACS) using a 100-micron nozzle on Sony SY3200 Cell Sorter.

#### 10x Genomics

Chromium Single Cell 3’ reagent Kits V2 were used for scRNA-seq library preparation following the manufacturer’s protocol.

#### Bulk RNA library preparation

RNA was prepared from dissected LL3 ovaries using QuiagenMicro Kit. The libraries were prepared with 5 ng total RNA input, using the NuGen Ovation RNA-Seq System V2, 7102-32, and the NuGen Ovation Ultralow System V2, 0344-32 kits, using the manufacturer’s protocol. The samples were sequenced in one lane of Hiseq4000, as paired end 150.

#### Sequencing

Single-cell RNAseq analysis was performed for 10X libraries sequenced on paired-end 26/98 Illumina HiSeq 4000 runs.

### QUANTIFICATION AND STATISTICAL ANALYSIS

#### 10x Genomics data preprocessing

Per-read per-sample FASTQ files were generated using the Illumina bcl2fastq Conversion software (v2.17) to convert BCL base call files outputted by the sequencing instrument into the FASTQ format.

The 10X Genomics analysis software, Cell Ranger (v1.3.1 for replicate 1 and v2.0.0 for replicate 2), specifically the “cellranger count” pipeline, was used to process the FASTQ files in order to align reads to the *Drosophila melanogaster* reference genome (dm6) (Santos et al., 2015) and generate gene-barcode expression matrices. The output of multiple samples from the “cellranger count” pipeline were aggregated using the “cellranger aggr” pipeline of Cell Ranger, normalizing the combined output to the same sequencing depth and recomputing the gene-barcode matrices and expression analysis accordingly for the aggregated data.

#### 10x Genomics data quality control

Seurat 2 package (Butler et al., 2018) was used for all scRNA-seq analysis. In brief, to remove low quality cells and potential doublets, we filtered out cells in which more than 5% of reads were from mitochondrial genes, and cells that express less than 500 genes. We had determined that germ cells express higher number of genes and UMIs than somatic cells, therefore, to filter out doublets, we set different filtering thresholds for somatic cells and germ cells. We identified germ cells by expression of five highly specific previously known and newly identified germ cell genes – *vas*, *ovo*, *bru1*, *AGO3* and *CG9926*. We filtered out cells germ cells in which we detected more than 90 000 UMIs and somatic cells with more than 60 000 UMIs.

#### scRNA-seq data analysis

The two scRNA-seq datasets were integrated (aligned) using Seurat v2 (Butler et al., 2018). We followed the Seurat v2 guidelines for identification of variable genes, dimensionality reduction and cell clustering. We used multiple resolution parameters (1.2 – 1.7) and obtained similar results (discussed in results). To find markers, we used Wilcox statistical test built in Seurat 2.

#### Transcriptional signatures

To compute transcriptional signatures for GC, SH, TF, CC, IC, FSCP and SW, we selected all the markers that are assigned to only one, two or three of these cell types.

#### GLAD analyses

We used the GLAD online tool (Hu et al., 2015) to determine if the marker genes for each cell type fall into particular gene categories. We used hypergeometric test to determine whether each gene category is significantly enriched in each cell type’s transcriptional signature.

#### Bulk RNA-seq data preprocessing

Per-read per-sample FASTQ files were generated using the Illumina bcl2fastq Conversion software (v2.20) to convert per-cycle BCL base call files outputted by the sequencing instrument into the FASTQ format. The alignment program, STAR (v2.4.5a) (Dobin et al., 2013), was used for mapping reads to the D. melanogaster reference genome dm6 (Santos et al., 2015) and the application FastQ Screen (v0.5.2) (Wingett and Andrews, 2018) was utilized to check for contaminants. The software, featureCounts (Subread package v1.4.6-p3) (Liao et al., 2013; 2014), was used to generate the matrix of read counts for annotated genomic features.

#### scRNA-seq and bulk RNA-seq correlation

The mean expression value was calculated for each gene among all cells in the scRNA seq datasets, transformed to log10 scale and plotted against log10 scaled counts of bulk-RNA seq data.

### DATA AND SOFTWARE AVAILABILITY

The scRNA-seq data has been deposited in GEO under accession code GEO: GSE131971.

---

**Figure S1.**
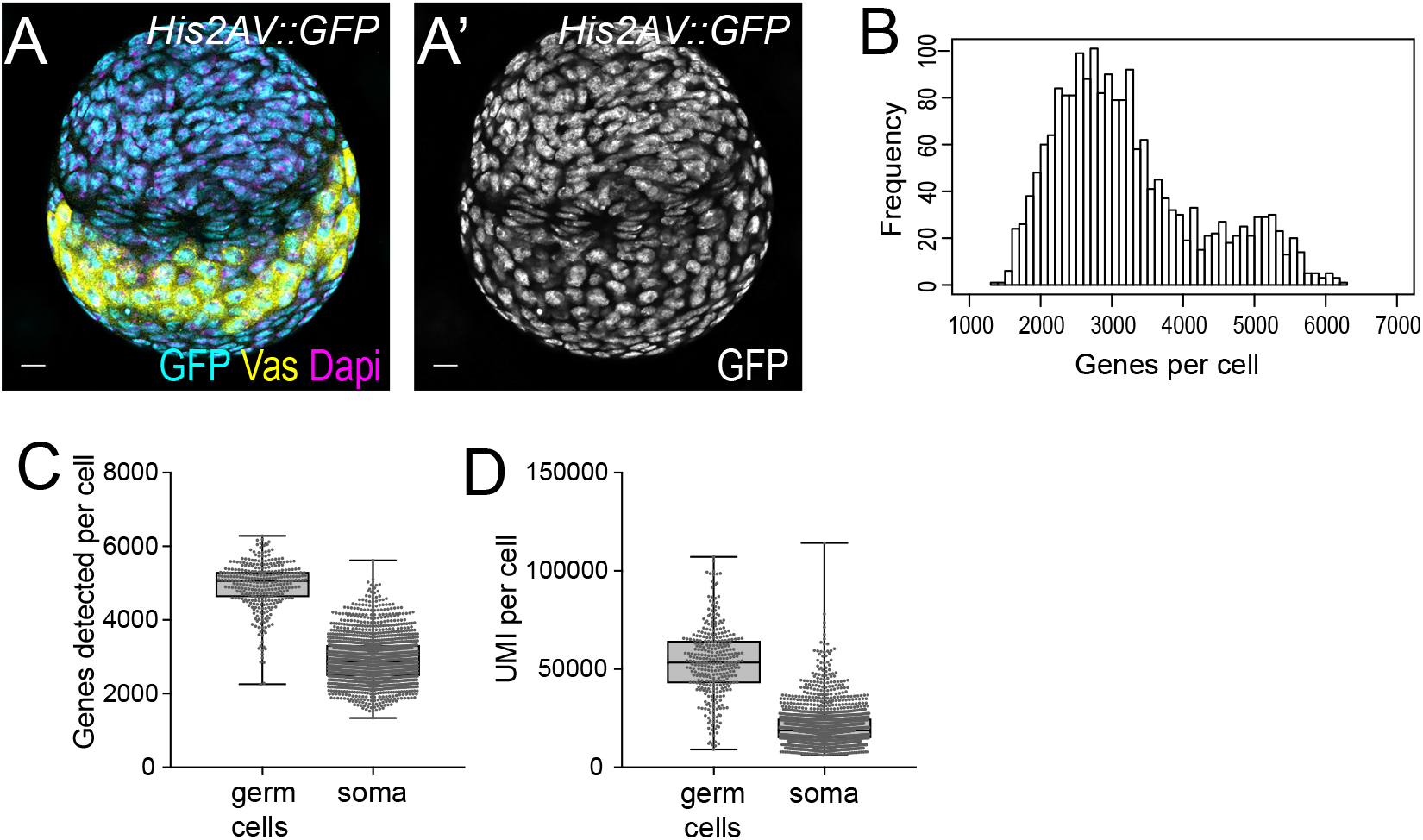
scRNA-seq experiment design and statistics. A – Immunofluorescence staining of developing ovaries at the time of dissections for scRNA-seq (LL3). GFP (cyan) labels all cell nuclei, Vas (yellow) labels GC cytoplasm, and Dapi (magenta) labels DNA. Scale bar - 10 m. B - Histogram of number of genes detected per cell. C – Box plots depicting number of genes detected in germ cells and somatic cells. D – Box plots of number of UMIs detected in germ cells and somatic cells. Whiskers show min and max value.

**Figure S2.**
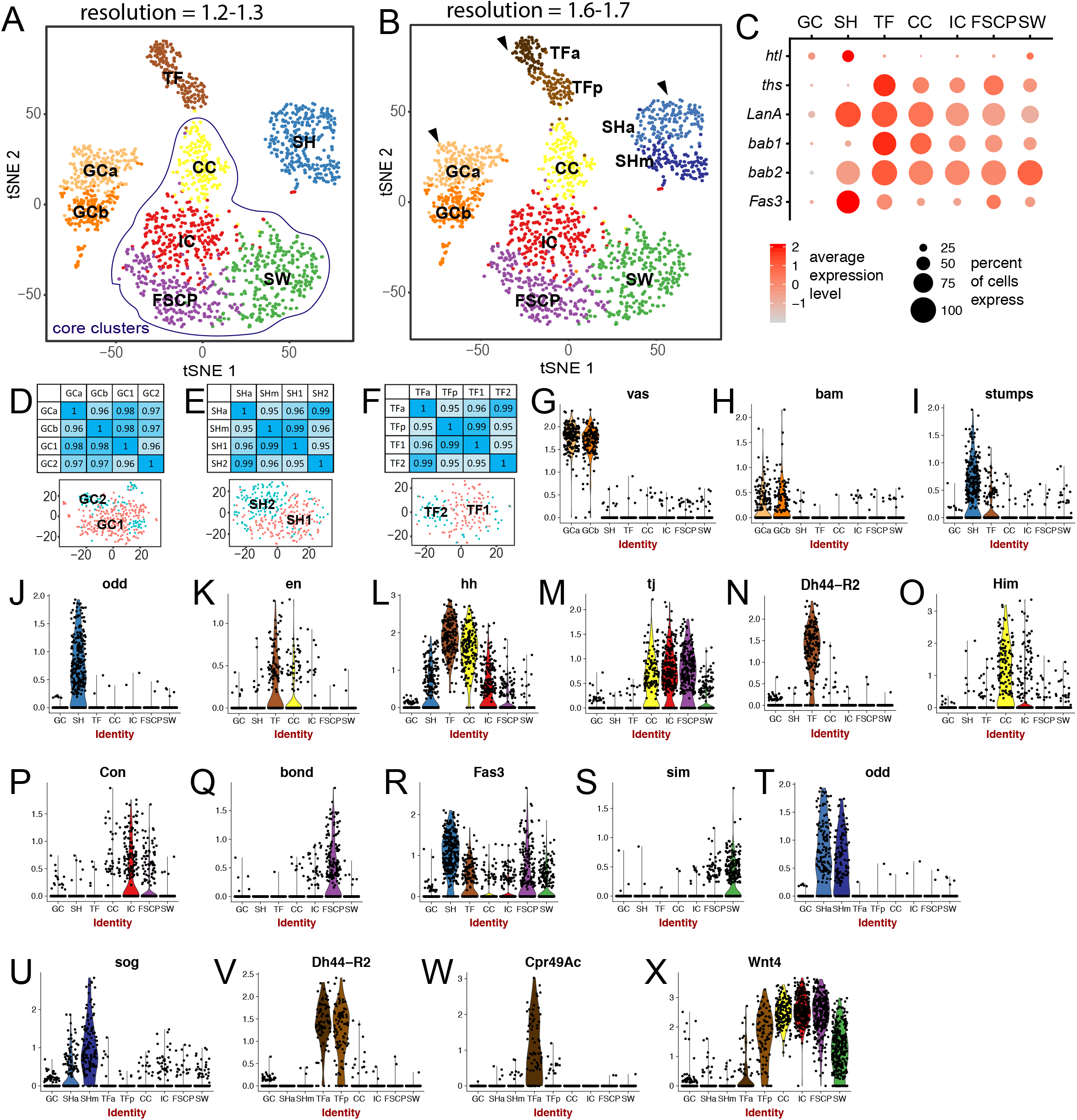

**Figure S3.**
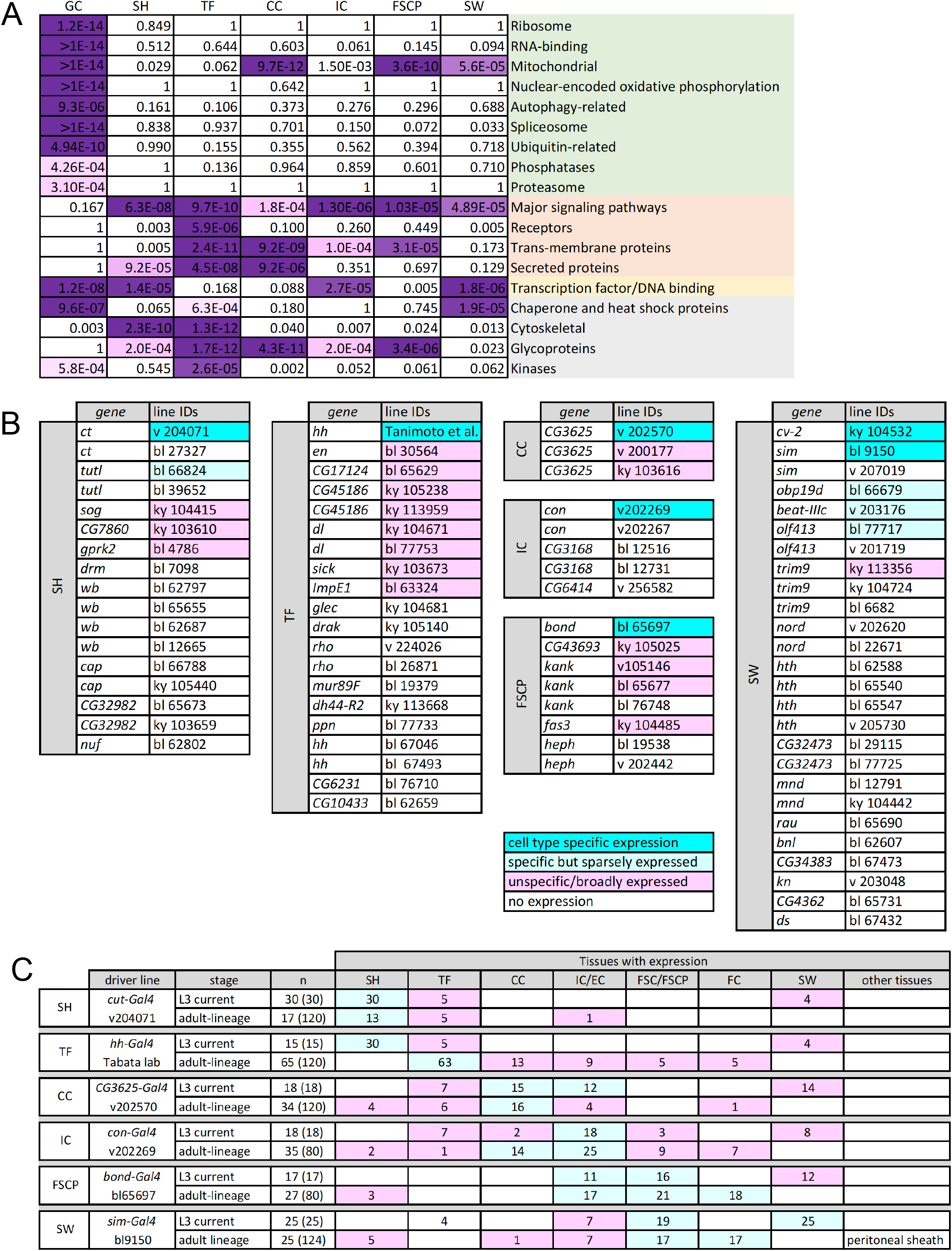

**Figure S4.**
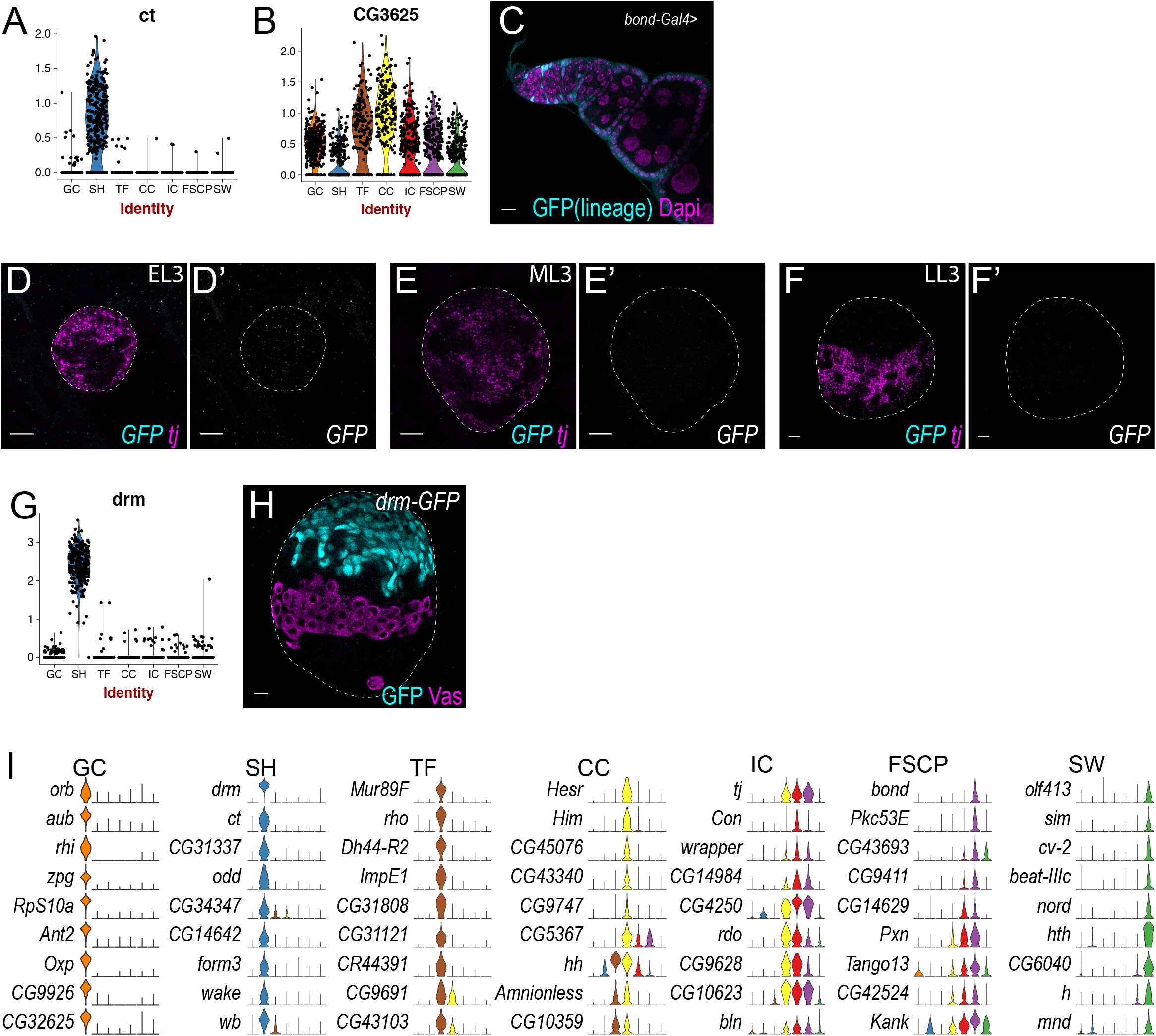
FSCPs give rise to FSCs and FCs in adult ovaries. A, B – Violin plots. A – ct is specifically expressed in SH. B – CG3625 has the highest expression in CCs, with lower expression in TFs, ICs and other cell types. Note - while CG3625 mRNA expression is rather broad, the CG3625-Gal4 line v202570, drives expression predominantly in CCs. C - Immunofluorescence of lineage tracing using bond-Gal4 (for FSCPs). GFP in cyan labels the lineage expression in FSCs and FCs. D-F - mRNA in situ hybridization using HCR at EL3 (D), ML3(E), and LL3 (F), tj (magenta) and GFP probes (cyan) as a negative control for Figure 4A-C. G – Violin plot. drm is specifically expressed in SH. H – Immunofluorescence staining of drm-GFP. GFP (cyan) labels SH, Vas (magenta) labels GCs. I – Violin plots visualizing expression patterns of nine most selectively expressed genes for each cell type.

