## Supplemental Note and Data for "A single cell atlas of the developing *Drosophila* ovary identifies follicle stem cell progenitors"

### SUPPLEMENTAL DATA

#### Supplemental Note.

##### 1. QC for scRNA-seq

In both scRNA-seq experiments, we detected a median of over 20,000 unique molecular identifiers (UMI, the barcodes, which label individual mRNAs) per cell that corresponded to more than 3,000 genes per cell was obtained. Among all cells in the ovary we detected 10,241 and 10,979 genes (of a total of about 17 thousand genes in *Drosophila*), respectively, in the two samples (Figure 1C). We used previously established quality control methods to exclude damaged cells and cell doublets. Cells with a high fraction of reads derived from genes encoded by the mitochondrial genome, an indicator of damaged cells, were excluded (Butler et al., 2018). A standard method for filtering out potential cell doublets, is to exclude transcriptomes in which significantly more genes than the median are detected. However, when we plotted cell distribution by number of genes expressed (nGene), we did not observe the expected normal distribution, but instead observed two peaks suggesting the existence of two cell populations that differ in the number of genes expressed (Figure S1B). Further investigation revealed that germ cell-specific genes were enriched in the cell population that expressed more genes and contained more transcripts (Figure S1CD). Therefore, we separated germ cells from somatic cells using a set of previously known and newly identified germ cell specific marker genes, and set different filtering thresholds for the number of expressed genes in the germline and soma populations (see Methods).

We detected more transcripts corresponding to higher number of genes in germ cells than in any somatic cell type. This finding could indicate that germ cells express higher number of genes. In support of this hypothesis, a recent study linked the increased number of expressed genes to a transcriptional scanning mechanism that may correct DNA damage during mouse spermatogenesis (Xia et al., 2018). Alternatively, since only a small fraction of all mRNAs in the cell is sequenced, higher RNA levels may indicate a higher RNA content. Consistent with this hypothesis, we detected higher number of UMIs in germ cells compared to somatic cells (Figure S1D). Indeed, germ cells are known for their reliance on post-transcriptional regulation (Slaidina and Lehmann, 2014) and mRNAs can be stored for days before they are translated (Tadros and Lipshitz, 2005).

#### 2. GC sub-clusters

When using higher resolution parameters for cell clustering, the GC cluster split into sub-clusters (Figure S2B, arrowheads). Closer analysis of gene expression patterns in the split germ cell cluster did not support the hypothesis that the two sub-clusters may correspond to GSCs and their differentiating progeny, since expression of the differentiation factor *bam* was the same in both sub-clusters (0.33 vs 0.42 in GCa and GCb, respectively, p-val = 0.11, Figure S2H). Furthermore, the sub-clusters did not present a clear signature, and similar clusters were not observed when GCs were re-clustered separately from the somatic cell types (Figure S2D). We conclude that this cluster split is unlikely to have biological significance but rather reflects an analytical error caused by unsupervised clustering of two cell populations, germ cells and somatic cells, with strikingly different expression levels and profiles.

#### KEY RESOURCES TABLE

| REAGENT or RESOURCE | SOURCE | IDENTIFIER |
|---|---|---|
| Antibodies | | |
| Chicken polyclonal anti GFP | Aves Labs Inc. | # GFP1020 |
| Rat monoclonal anti RFP | Chromotek | # 5F8 |
| Guinea pig polyclonal anti Tj | Dorothea Godt Lab | |
| Mouse monoclonal anti Fas3 | DSHB | #7G10 |
| Rabbit anti Vasa | Lehmann lab | |
| Chemicals, Peptides, and Recombinant Proteins | | |
| DPBS (no calcium, no magnesium) | Thermo Fisher Scientific | Cat#14190144 |
| Trypsin 1:250 | Thermo Fisher Scientific | Cat#27250018 |
| Type I Collagenase | Invitrogen (Thermo Fisher Scientific) | Cat#17018029 |
| Chromium Single Cell 3' Library & Gel Bead Kit v2 | 10x Genomics | Cat#PN-120237 |
| Chromium Single Cell A Chip Kit | | Cat#PN-1000009 |
| Critical Commercial Assays | | |
| In Situ HCR v3.0 mRNA Imaging Kit (HCR probe sets, HCR amplifiers, HCR buffers (Probe hybridization buffer, Probe wash buffer, Amplification buffer) | Molecular Instruments | Custom |
| Experimental Models: Organisms/Strains | | |
| *Drosophila melanogaster : $w^{1118}$* | Lehmann lab stock | |
| *Drosophila melanogaster : His2AV::GFP :* $w^{1118}$; P{His2Av$^{T:Avic\backslash GFP-S65T}$}62A | (Clarkson and Saint, 1999) | BDSC # 5941 |
| *Drosophila melanogaster : drm-GFP : 2XTY1-T2A-SGFP-NLS-3XFLAG* | (Sarov et al., 2016) | VDRC # 318404 |

| | | |
|---|---|---|
| *Drosophila melanogaster : GTRACE :* w[*]; P{w[+mC]=UAS-RedStinger}4, P{w[+mC]=UAS-FLP.D}JD1, P{w[+mC]=Ubi-p63E(FRT.STOP)Stinger}9F6/CyO | (Evans et al., 2009) | BDSC # 28280 |
| *Drosophila melanogaster : GTRACE :* w[*]; P{w[+mC]=UAS-RedStinger}6, P{w[+mC]=UAS-FLP.Exel}3, P{w[+mC]=Ubi-p63E(FRT.STOP)Stinger}15F2 | (Evans et al., 2009) | BDSC # 28281 |
| *Drosophila melanogaster : cut*-Gal4 : VT058382.GAL4@attP2 | (Tirian and Dickson, 2017) | VDRC # 204071 |
| *Drosophila melanogaster : CG3625*-Gal4 : VT000131.GAL4@attP2 | (Tirian and Dickson, 2017) | VDRC # 202570 |
| *Drosophila melanogaster : Con*-Gal4 : VT025803.GAL4@attP2 | (Tirian and Dickson, 2017) | VDRC # 202269 |
| *Drosophila melanogaster : bond-Gal4 :* PBac{IT.GAL4}bond[1385-G4] | (Gohl et al., 2011) | BDSC # 65697 |
| *Drosophila melanogaster : sim-Gal4 :* w[*]; P{w[+mC]=GAL4-sim.3.7}2/CyO; P{w[+mC]=GAL4-sim.3.7}3 | (Shen et al., 2013) | BDSC # 9150 |
| *Drosophila melanogaster : hh*-Gal4 | (Tanimoto et al., 2000) | |
| *Drosophila melanogaster : UAS-rpr :* w[1118]; P{w[+mC]=UAS-rpr.C}14 | (Aplin and Kaufman, 1997) | BDSC # 5824 |
| Software and Algorithms | | |
| Cell Ranger v1.3.1, v2.0.0 | 10x Genomics | |
| Seurat 2 | (Butler et al., 2018) | |
| STAR v2.4.5a | (Dobin et al., 2013) | |
| featureCounts (Subread package v0.5.2) | (Liao et al., 2013; 2014) | |
| FastQ Screen(c0.5.2) | (Wingett and Andrews, 2018) | |
| Fiji | (Schindelin et al., 2012) | |
